# Light-entrained and brain-tuned circadian circuits regulate ILC3 and gut homeostasis

**DOI:** 10.1101/723932

**Authors:** Cristina Godinho-Silva, Rita G. Domingues, Miguel Rendas, Bruno Raposo, Hélder Ribeiro, Joaquim Alves da Silva, Ana Vieira, Rui M. Costa, Nuno L. Barbosa-Morais, Tânia Carvalho, Henrique Veiga-Fernandes

## Abstract

Group 3 innate lymphoid cells (ILC3) are major regulators of inflammation, infection, microbiota composition and metabolism^1^. ILC3 and neuronal cells were shown to interact at discrete mucosal locations to steer mucosal defence^2,3^. Nevertheless, whether neuroimmune circuits operate at an organismal level, integrating extrinsic environmental signals to orchestrate ILC3 responses remains elusive. Here we show that light-entrained and brain-tuned circadian circuits regulate enteric ILC3, intestinal homeostasis, gut defence and the host lipid metabolism. We found that enteric ILC3 display circadian expression of clock genes and ILC3-related transcription factors. ILC3-autonomous ablation of the circadian regulator *Arntl* led to disrupted gut ILC3 homeostasis, impaired epithelial reactivity, deregulated microbiome, increased susceptibility to bowel infection and disrupted lipid metabolism. Loss of ILC3-intrinsic *Arntl* shaped the gut postcode receptors of ILC3. Strikingly, light-dark cycles, feeding rhythms and microbial cues differentially regulated ILC3 clocks, with light signals as major entraining cues of ILC3. Accordingly, surgical- and genetically-induced deregulation of brain rhythmicity led to disrupted circadian ILC3 oscillations, deregulated microbiome and altered lipid metabolism. Our work reveals a circadian circuitry that translates environmental light cues into enteric ILC3, shaping intestinal health, metabolism and organismal homeostasis.

---

Group 3 innate lymphoid cells (ILC3) were shown to be part of discrete mucosal neuroimmune cell units^2-5^, raising the hypothesis that ILC3 may also integrate systemic neuroimmune circuits to regulate tissue integrity and organismic homeostasis. Circadian rhythms rely on local and systemic cues to coordinate mammalian physiology and are genetically encoded by molecular clocks that allow organisms to anticipate and adapt to extrinsic environmental changes^6,7^. The circadian clock machinery consists of an autoregulatory network of feedback loops primarily driven by the activators ARNTL and CLOCK and the repressors PER1-3 and CRY1-2, amongst others^6,7^.

Analysis of intestinal ILC subsets and their bone marrow (BM) progenitors revealed that mature ILC3 express high levels of circadian clock genes (Fig.1a-c and Extended Data Fig.1a-d). Importantly, ILC3 displayed a circadian pattern of *Per1*^Venus^ expression (Fig.1b) and transcriptional analysis of ILC3 revealed circadian expression of master clock regulators and ILC3-related transcription factors (Fig.1c). To explore if ILC3 are regulated in a circadian manner, we investigated whether intestinal ILC3 require intrinsic clock signals. Thus, we interfered with the expression of the master circadian activator *Arntl. Arntl*^fl^ mice were bred to *Vav1*^Cre^ mice, allowing for conditional deletion of *Arntl* in all hematopoietic cells (*Arntl*^ΔVav1^ mice). While *Arntl*^ΔVav1^ mice displayed normal numbers of intestinal NK cells and enteric groups 1 and 2 ILC, gut ILC3 were severely and selectively reduced when compared to their wild type littermate controls (Fig.1d,e and Extended Data Fig.2a,b). To more precisely define ILC3-intrinsic effects, we performed mixed BM chimeras transferring *Arntl* competent (*Arntl*^fl^) or deficient (*Arntl*^ΔVav1^) BM against a third-part wild type competitor into alymphoid hosts (Fig.1f). Analysis of such chimeras confirmed a cell-autonomous circadian regulation of ILC3, while their innate and adaptive counterparts were unperturbed (Fig.1g and Extended Data Fig.2c).

To explore the functional impact of ILC3-intrisic circadian signals, we deleted *Arntl* in RORγt expressing cells by breeding *Rorgt*^Cre^ to *Arntl*^fl^ mice (*Arntl*^ΔRorgt^ mice). When compared to their wild type littermate controls, *Arntl*^ΔRorgt^ mice revealed selective reduction of ILC3 subsets and IL-17- and IL-22-producing ILC3 (Fig.2a,b and Extended Data Fig.3a-j). Noteworthy, independent deletion of *Nr1d1* also perturbed enteric ILC3 subsets, further supporting a role of the clock machinery in ILC3 (Extended Data Fig.4a-e). ILC3 were shown to regulate epithelial reactivity gene expression and microbial composition^1^. Analysis of *Arntl*^fl^ and *Arntl*^ΔRorgt^ mice revealed a profound reduction of reactivity genes in the *Arntl*^ΔRorgt^ intestinal epithelium; notably, *Reg3b, Reg3g, Muc3* and *Muc13* were consistently reduced in *Arntl* deficient mice (Fig.2c). Furthermore, *Arntl*^ΔRorgt^ mice displayed altered diurnal patterns of Proteobacteria and Bacteroidetes (Fig.2d and Extended Data Fig.3j). To interrogate whether disruption of ILC3-intrinsic ARNTL impacted enteric defence, we tested how *Arntl*^ΔRorgt^ mice responded to intestinal infection. To this end, *Arntl*^ΔRorgt^ mice were bred to *Rag1*^-/-^ mice to exclude putative T cell effects (Extended Data Fig.3g-i). *Rag1*^-/-^*.Arntl*^ΔRorgt^ mice were infected with the attaching and effacing bacteria *Citrobacter rodentium*^2^. When compared to their wild type littermate controls, *Rag1*^-/-^*.Arntl*^ΔRorgt^ mice had marked gut inflammation, reduced IL-22 producing ILC3, increased *C. rodentium* infection and bacterial translocation, reduced epithelial reactivity genes, increased weight loss and reduced survival (Fig.2e-j and Extended Data Fig.5a-j). These results indicate that cell-intrinsic circadian signals selectively control intestinal ILC3 and shape gut epithelial reactivity, microbial communities and enteric defence. Previous studies indicated that ILC3 regulate host lipid metabolism^8^. When compared to their wild type littermate controls, the *Arntl*^ΔRorgt^ epithelium revealed a marked increase in mRNA coding for key lipid epithelial transporters, including *Fabp1, Fabp2, Scd1*, C*d36* and *Apoe* (Fig.2k). Accordingly, these changes associated with increased gonadal and subcutaneous fat accumulation in *Arntl*^ΔRorgt^ mice when compared to their wild type littermate controls (Fig.2l and Extended Data Fig.5k-n). Thus, ILC3-intrinsic circadian signals shape epithelial lipid transport and body fat composition.

**Figure 1.**
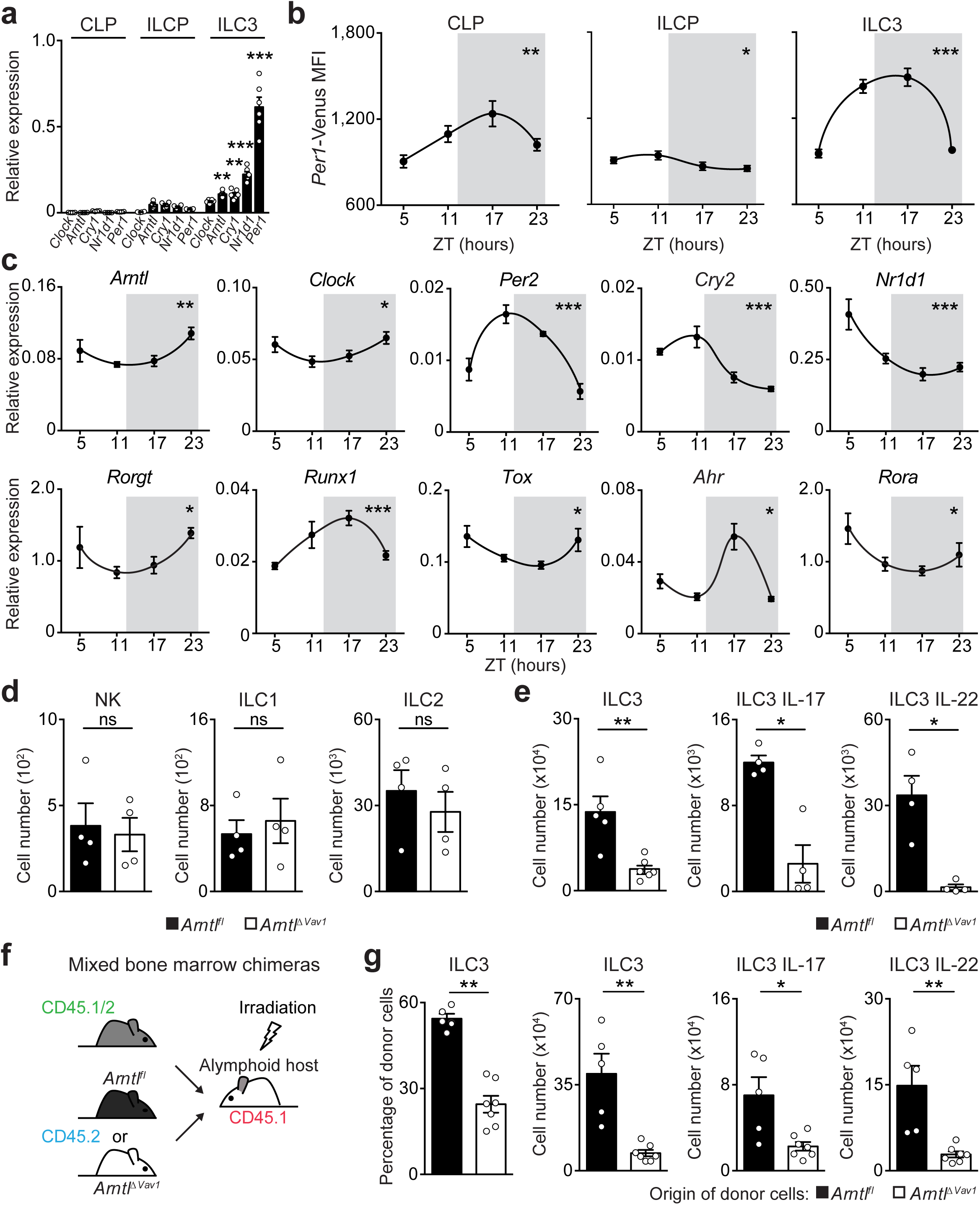
Intestinal ILC3 are controlled in a circadian manner. **a**, Common lymphoid progenitor (CLP); innate lymphoid cell progenitors (ILCP); intestinal ILC3. CLP and ILCP n=4; ILC3 n=6. **b**, *Per1*^Venus^ MFI. CLP and ILCP n=6; ILC3 n=4. **c**, enteric ILC3. n=5. **d**, Intestinal ILC subsets. n=4. **e**, Gut ILC3. n=4. **f, g**, Mixed BM chimeras. *Arntl*^fl^ n=5, *Arntl1*^ΔVav1^ n=7. (b-d) White/Grey: light/dark period. Data are representative of 3 independent experiments. n represents biologically independent samples (a,c) or animals (b,d-g). Mean and error bars: s.e.m. (a) two-way ANOVA and Tukey’s test; (b,c) Cosinor analysis; (d,e,g) two-tailed Mann-Whitney U test. *P<0.05; **P<0.01; ***P<0.001; ns not significant.

**Figure 2.**
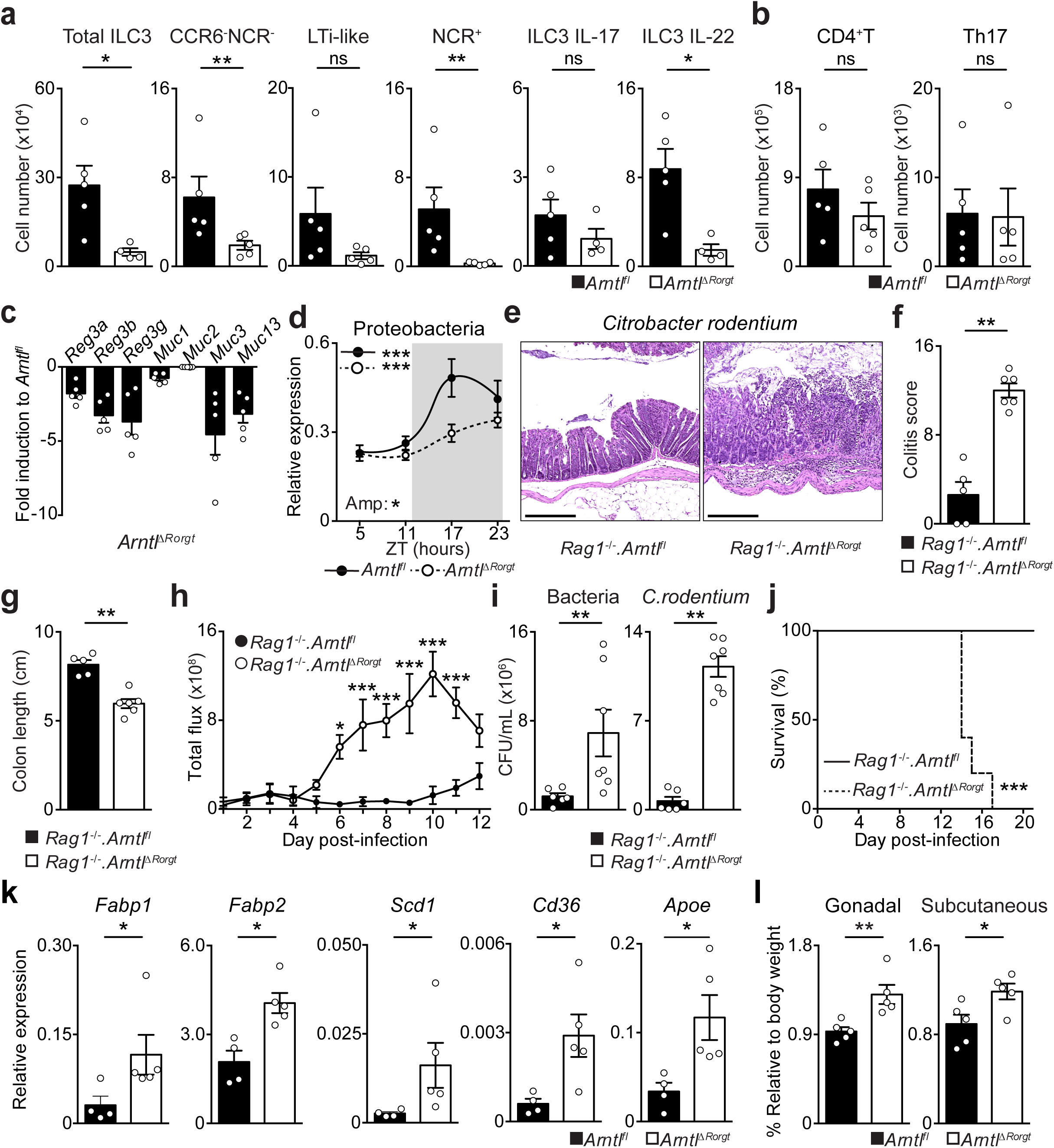
ILC3-intrinsic *Arntl* regulates gut homeostasis and defence. **a**, Enteric ILC3. n=4. **b**, Gut T helper cells. n=5. **c**, Epithelial reactivity genes. n=5 **d**, Proteobacteria. *Arntl*^fl^ n=5; *Arntl*^ΔRorgt^ n=6. **e**, Colon of *C. rodentium* infected mice. n=5. **f**, Colitis score. n=5. **g**, Colon length. n=5. **h**, Infection burden. *Rag1*^-/-^*.Arntl*^fl^ n=6, *Rag1*^-/-^*.Arntl*^ΔRorgt^ n=7. **i**, Bacterial translocation to the spleen. *Rag1*^-/-^*.Arntl*^fl^ n=6, *Rag1*^-/-^ *.Arntl*^ΔRorgt^ n=7. **j**, Survival. n=5; **k**, Epithelial lipid transporter genes. *Arntl*^fl^ n=4; *Arntl*^ΔRorgt^ n=5. **l**, Adipose tissue. n=5. (d) White/Grey: light/dark period. Scale bars: 250µm. Data are representative of at least 3 independent experiments. (a-l) n represents biologically independent animals. Mean and error bars: s.e.m. (a,b,f,g,i,k) two-tailed Mann-Whitney U test; (d) Cosinor analysis; (h) two-way ANOVA and Sidak’s test; (j) Log-rank test; (l) two-tailed unpaired Student t-test. *P<0.05; **P<0.01; ***P<0.001; ns not significant.

To further examine how cell-intrinsic *Arntl* controls intestinal ILC3 homeostasis, initially we interrogated the diurnal oscillations of the ILC3 clock machinery. When compared to their wild type littermate controls, *Arntl*^ΔRorgt^ ILC3 displayed a disrupted diurnal pattern of activator and repressor circadian genes (Fig.3a). Sequentially, we employed genome-wide transcriptional profiling of *Arntl* sufficient and deficient ILC3 to interrogate the impact of a deregulated circadian machinery. Diurnal analysis of the genetic signature associated with ILC3 identity^1^ demonstrated that the vast majority of those genes were unperturbed in *Arntl* deficient ILC3, suggesting that ARNTL is dispensable to ILC3-lineage commitment (Fig.3b and Extended Data Fig.6a-c). To test this hypothesis, we first interrogated the impact of *Arntl* ablation in ILC3 progenitors. *Arntl*^ΔVav1^ mice had unperturbed numbers of common lymphoid progenitors (CLP) and innate lymphoid cell progenitors (ILCP) (Fig.3c and Extended Data Fig.6d). Sequentially, we analysed the impact of *Arntl* ablation in ILC3 residing in other organs. Compared to their littermate controls, *Arntl*^ΔRorgt^ mice revealed normal numbers of ILC3 in the spleen, lung and blood, which was in large contrast to their pronounced reduction in the intestine (Fig.2a; Fig.3d,e and Extended Data 6e). Noteworthy, enteric *Arntl*^ΔRorgt^ ILC3 displayed unperturbed proliferation and apoptosis-related genetic signatures (Extended Data Fig.6b,c), suggesting that *Arntl*^ΔRorgt^ ILC3 may have altered intestinal traffic^9^. When compared to their wild type littermate controls, *Arntl*^ΔRorgt^ ILC3 had a marked reduction of essential receptors for intestinal lamina propria homing and accumulated in mesenteric lymph-nodes (Extended Data Fig.6f)^9^. Notably, the integrin and chemokine receptors CCR9, α4β7 and CXCR4 were selectively and hierarchically affected in *Arntl*^ΔRorgt^ ILC3 (Fig.3f-h and Extended Data Fig.6g-m). To examine whether ARNTL could directly regulate *Ccr9* expression we performed chromatin immunoprecipitation. Binding of ARNTL to the *Ccr9* locus of ILC3 followed a diurnal pattern, with increased binding at ZT5 (Fig.3i). Thus, ARNTL can directly contribute to *Ccr9* expression in ILC3, although additional factors may also regulate this gene. In conclusion, while a fully operational ILC3-intrinsic circadian machinery is dispensable to ILC3-lineage commitment and development, cell-intrinsic clock signals are required for a functional ILC3 gut receptor postcode.

**Figure 3.**
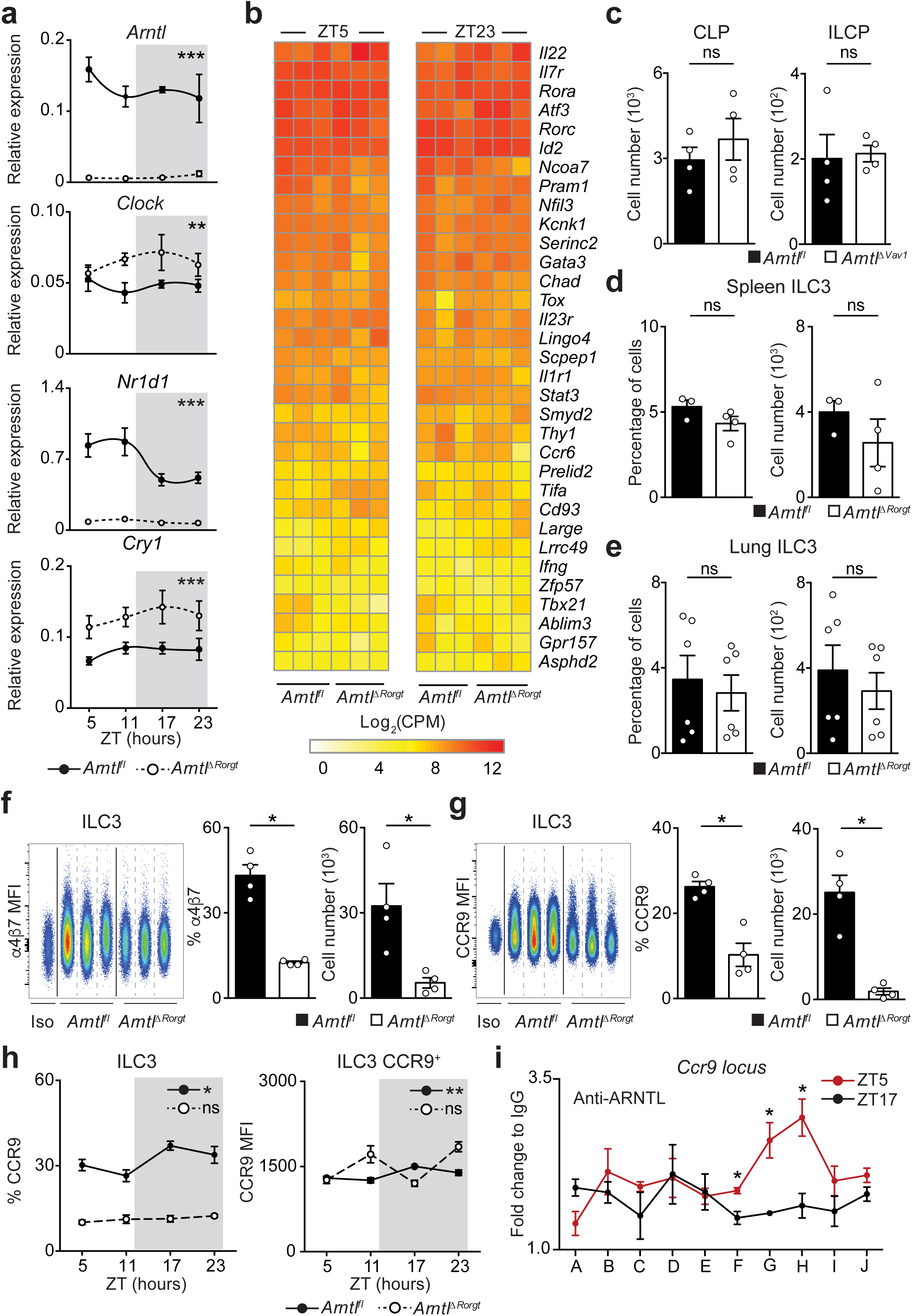
ILC3-intrinsic circadian signals regulate an enteric receptor postcode. **a**, Enteric ILC3. n=3. **b**, RNA-seq. Gut ILC3. n=3. **c**, CLP and ILCP. n=4. **d, e**, ILC3. (d) spleen, n=3; (e) lung n=6. **f, g**, Gut ILC3. n=4. **h**, intestinal ILC3. n=4. **i**, ChIP analysis in enteric ILC3. n=3. Data are representative of 3 independent experiments. n represents biologically independent animals (a,c-h) or samples (b,i). (i) A-J putative ARNTL DNA binding sites. (a,h) White/Grey: light/dark period. Mean and error bars: s.e.m. (a) two-way ANOVA; (c-g) two-tailed Mann-Whitney U test; (h) Cosinor analysis; (i) two-tailed unpaired Student t-test. *P<0.05; **P<0.01; ***P<0.001; ns not significant.

Circadian rhythms allow organisms to adapt to extrinsic environmental changes. Microbial cues were shown to impact the rhythms of intestinal cells^10,11^, while feeding regimens are major circadian entraining cues of peripheral organs, such as the liver^12^. In order to define the environmental cues that entrain circadian oscillations of ILC3, initially we investigated whether microbial cues impact the oscillations of ILC3. *Per1*^Venus^ reporter mice treated with antibiotics displayed an unperturbed circadian oscillatory amplitude, while exhibiting a minute shift of their acrophase (Fig.4a). Sequentially, we tested whether feeding regimens, known to be major entraining cues of liver, pancreas, kidney, and heart oscillations^12^, impact ILC3 rhythms. To this end, we restricted food access to a 12 hours interval and compared *Per1*^Venus^ oscillations to those observed in mice with inverted feeding regimens^12^. Inverted feeding had a small impact in the amplitude of ILC3 oscillations but did not invert the acrophase of ILC3 (Fig.4b and Extended Data Fig.7a), which was in large contrast with the full inversion of the acrophase of hepatocytes (Extended Data Fig.7b)^12^. Since these local intestinal cues could not invert the acrophase of ILC3, we hypothesise that light-dark cycles are major regulators of enteric ILC3 oscillations^6^. To test this hypothesis, we placed *Per1*^Venus^ mice in light-tight cabinets on two opposing 12 hours light-dark cycles. Strikingly, inversion of light-dark cycles had a profound impact in the circadian oscillations of ILC3 (Fig.4c). Notably, and in contrast to microbiota and feeding regimens, light cycles fully inverted the acrophase of *Per1*^Venus^ oscillations in ILC3 (Fig.4c and Extended Data Fig.7c). Furthermore, light-dark cycles entrained ILC3 oscillations as revealed by their maintenance upon removal of light (constant darkness) (Fig.4d and Extended Data Fig.7d), formally supporting that light is a major environmental entraining signal of ILC3 intrinsic oscillations. Together, these data indicate that ILC3 integrate systemic and local cues hierarchically; while microbiota and feeding regimens locally adjust the ILC3 clock, light-dark cycles are major entraining cues of ILC3, fully setting and entraining their intrinsic oscillatory clock.

The suprachiasmatic nuclei (SCN) in the hypothalamus are main integrators of light signals^6^, suggesting that brain cues may regulate ILC3. To assess the impact of the master circadian pacemaker in ILC3, while excluding confounding light-induced, SCN-independent effects^13,14^, we performed SCN ablation by electrolytic lesion in *Per1*^Venus^ mice using stereotaxic brain surgery^15^. Strikingly, while sham operated mice displayed circadian *Per1*^Venus^ oscillation in ILC3, their counterparts from SCN ablated mice fully lost the circadian rhythmicity of *Per1*^Venus^ and other circadian genes (Fig.4e,f and Extended Data Fig.8a-d). Since electrolytic lesions of the SCN may cause scission of afferent and efferent fibres in the SCN, we further confirmed that brain SCN-derived cues control ILC3 by the genetic ablation of *Arntl* in the SCN^14^. *Arntl*^fl^ mice were bred to *Camk2a*^Cre^ mice allowing for a forebrain/SCN-specific deletion of *Arntl* (*Arntl*^ΔCamk2a^)^14^. When compared to their control counterparts, ILC3 from *Arntl*^ΔCamk2a^ mice revealed severe arrhythmicity of circadian regulatory genes and of the enteric postcode molecule CCR9 (Fig.4g,h and Extended Data Fig.9a-f). Importantly, *Arntl*^ΔCamk2a^ mice further exhibited altered epithelial reactivity genes and perturbed microbial communities, notably Proteobacteria and Bacteroidetes (Fig.4i,j and Extended Data Fig.9g-i). Finally, the *Arntl*^ΔCamk2a^ intestinal epithelium showed disrupted circadian expression of lipid epithelial transporters, and these changes associated with increased gonadal and subcutaneous fat accumulation (Fig. 4k,l). Taken together, these data indicate that light-entrained and brain-tuned circuits regulate enteric ILC3, controlling microbial communities, lipid metabolism and body composition.

**Figure 4.**
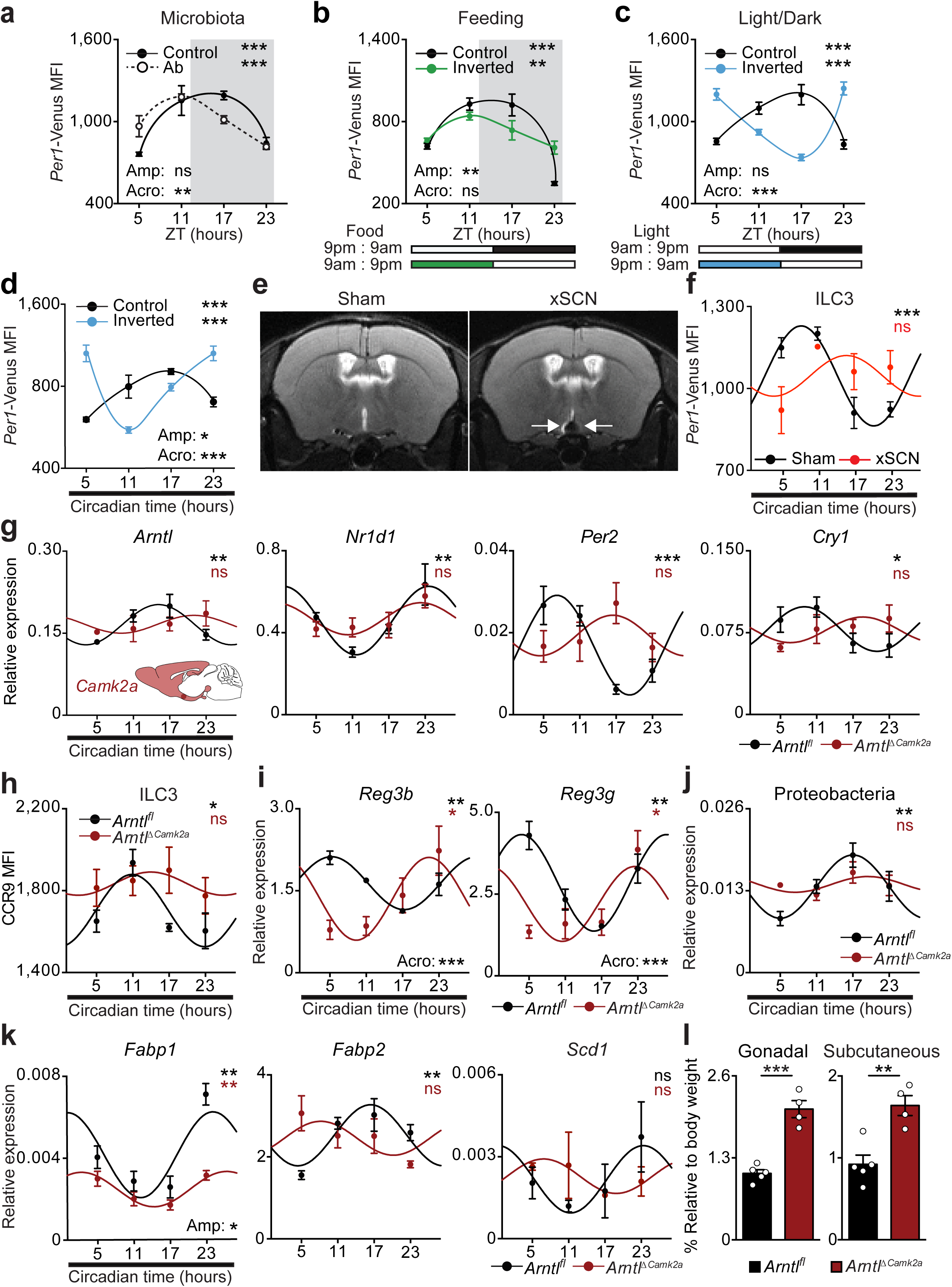
Light-entrained and brain-tuned cues shape intestinal ILC3. **a**, Antibiotic treatment. *Per1*^Venus^ in gut ILC3. n=3. **b**, Restricted and inverted feeding. *Per1*^Venus^ in gut ILC3. n=3. **c**, Opposing light-dark cycles. *Per1*^Venus^ in gut ILC3. n=3. **d**, Opposing light-dark cycles followed by constant darkness. *Per1*^Venus^ in gut ILC3. n=3. **e**, Magnetic resonance imaging of sham and SCN ablated (xSCN) mice. n=11. **f, g**, Enteric ILC3. n=3. **h**, Gut ILC3. n=3. **i**, Epithelial reactivity genes. n=3. **j**, Proteobacteria. n=4. **k**, Epithelial lipid transporter genes. n=3. **l**, Adipose tissue. *Arntl*^fl^ n=5, *Arntl*^ΔCamk2a^ n=4. (a,b) White/Grey: light/dark period. Mean and error bars: s.e.m.. (a-l) n represents biologically independent animals. (a-d;f-k) Cosinor analysis; (f-k) Cosine fitted curves; amplitude (Amp) and acrophase (Acro) were extracted from the Cosinor model. (l) two-tailed unpaired Student t-test. *P<0.05; **P<0.01; ***P<0.001; ns not significant.

Deciphering the mechanisms by which neuroimmune circuits operate to integrate extrinsic and systemic signals is critical to understand tissue and organ homeostasis. We found that light cues are major extrinsic entraining cues of ILC3 circadian rhythms and surgical- and genetically-induced deregulation of brain rhythmicity resulted in altered ILC3 regulation. In turn, the ILC3-intrinsic circadian machinery controlled the gut receptor postcode of ILC3, shaping enteric ILC3 and host homeostasis.

Our data reveal that ILC3 display diurnal oscillations that are genetically encoded, cell-autonomous and entrained by light cues. While microbiota and feeding regimens could locally induce small adjustments to ILC3 oscillations, light-dark cycles were major entraining cues of the ILC3 circadian clock. Whether the effects of photonic signals on ILC3 are immediate or rely on other peripheral clocks remains to be elucidated^16,17^. Nevertheless, cell-intrinsic ablation of important endocrine and peripheral neural signals in ILC3 did not affect gut ILC3 numbers (Extended Data Fig.10a-i). Our work indicates that ILC3 integrate local and systemic entraining cues in a distinct hierarchical manner, establishing an organismal circuitry that is an essential link between the extrinsic environment, enteric ILC3, gut defence, lipid metabolism and host homeostasis (Extended Data Fig.10j).

Previous studies demonstrated that ILC integrate tissue microenvironmental signals, including cytokines, micronutrients and neuroregulators^3,4,18,19^. Here we show that ILC3 have a cell-intrinsic circadian clock that integrates extrinsic light-entrained and brain-tuned signals. Coupling light cues to ILC3 circadian regulation may have ensured efficient and integrated multi-system anticipatory responses to environmental changes. Notably, the regulation of ILC3 activity by systemic circadian circuits may have been selected to maximize metabolic homeostasis, gut defence and efficient symbiosis with commensals that have been evolution partners of mammals. Finally, our current data may also contribute to a better understanding of how circadian disruptions in humans associate with metabolic diseases, bowel inflammatory conditions and cancer^20^.

## Acknowledgements

We thank the Vivarium, Flow Cytometry, Histology, Molecular Biology and Hardware platforms at the Champalimaud Centre for the Unknown. We thank the Congento infrastructure for genetic model organisms. We thank Roksana Pirzgalka, Roel Klein Wolterink, Sara Correia, Inês Godinho, Filipa Cardoso, Bethania Garcia Cassani and Kristin Fischer for technical help and discussions; Catherine French for helping in behaviour analysis; Noam Shemesh, Teresa Serradas Duarte and Daniel Nunes for MRI imaging. Artur Silva for technical help with hardware; Filipa Rijo-Ferreira and Leopoldo Petreanu for helpful discussions. Pedro Faísca for pathology scoring. C.G.-S., R.G.D and M.R were supported by Fundação para a Ciência e Tecnologia (FCT), Portugal. N.L.B.-M. is supported by FCT, Portugal, and European Molecular Biology Organisation (EMBO). H.V.-F. by ERC (647274), EU, The Paul G. Allen Frontiers Group, US, and FCT, Portugal.

## Author contribution

C.G.-S. and R.G.D designed, performed and analysed the experiments in Fig.1-4; and Extended data Fig.1-10. M.R. performed circadian analysis. M.R. and H.R provided technical assistance in Fig.2d-j and Extended Data Fig.5. M.R. provided technical assistance in Fig.3b; Extended Data Fig.6b,e. B.R. provided technical assistance in Fig.1a-d; Fig.3a,h,i; Fig.4j-k; Extended Data Fig.1, Extended Data Fig.3f,j; Extended Data Fig.6a,k,l and Extended Data Fig.8d; Extended Data Fig.9e-h. H.R. managed the animal colony. J.A.S. and R.M.C. helped designing the experiments in Fig.4e and Extended Data Fig.8a. A.V. provided technical assistance with flow cytometry. N.L.B.-M. analysed the experiments in Fig.3b and Extended Data Fig.6b. T.C. analysed the experiments in Fig.2e,f and Extended Data Fig.5a-c and Extended Data Fig.8d. H.V.-F. supervised the work, planned the experiments and wrote the manuscript.

## Author information

The authors declare no competing financial interests.

## Data availability

Source data for quantifications shown in all graphs plotted in figures and extended data figures are available in the online version of the paper. The data sets generated in this study are also available from the corresponding author upon reasonable request. RNA–seq datasets analysed are publicly available in Gene Expression Omnibus repository with the accession number GSE135235.

## Methods

### Mice

Nod/Scid/Gamma (NSG) were purchased from the Jackson Laboratories. C57BL/6J Ly5.1 were purchased from the Jackson Laboratories and bred with C57BL/6J in order to obtain C57BL/6 Ly5.1/Ly5.2 (CD45.1/CD45.2). *Rag1*^-/- 21^, *Rag2*^-/-^ *Il2rg*^-/- 22,23^, *Vav1*^Cre 24^, *Rorgt*^Cre 25^, *Camk2a*^Cre 26^, *Il7ra*^Cre 27^, *Per1*^Venus 28^, *Ret*^GFP 29^, *Rosa26*^RFP 30^, *Nr1d1*^-/- 31^, *Arntl*^fl 32^, *Nr3c1*^fl 33^ and *Adrb2*^fl 34^. All mouse lines were on a full C57BL/6J background. All lines were bred and maintained at Champalimaud Centre for the Unknown (CCU) animal facility under specific pathogen free conditions. 8-14 weeks old males and females were used in this study, unless stated otherwise. Sex and age matched mice were used for small intestine epithelium lipid transporters analysis and white adipose tissue quantification. Mice were maintained in 12h light-dark cycles, with *ad libitum* access to food and water, if not specified otherwise. For light inversion experiments mice were housed in ventilated, light-tight cabinets on defined 12h light-dark cycles (Ternox). *Camk2a*^Cre^.*Arntl*^*fl*^ (*Arntl*^ΔCamk2a^) and their WT littermate controls were maintained in constant darkness as previously described^14^. Mice were systematically compared with co-housed littermate controls unless stated otherwise. Power analysis was performed to estimate the number of experimental mice. All animal experiments were approved by national and local Institutional Review Boards (IRB), respectively, Direção Geral de Veterinária and CCU ethical committees. Randomisation and blinding were not used unless stated otherwise.

### Cell isolation

Isolation of small intestine and colonic lamina propria cells was previously described^2^. Briefly, intestines and colons were thoroughly rinsed with cold PBS1X, Peyer patches were removed from the small intestine, and intestines and colons were cut in 1cm pieces, and shaken for 30min in PBS containing 2% FBS, 1% HEPES and 5mM EDTA to remove intraepithelial and epithelial cells. Intestines and colons were then digested with collagenase D (0.5mg/mL; Roche) and DNase I (20U/mL; Roche) in complete RPMI for 30min at 37°C, under gentle agitation. Sequentially cells were passed through a 100μM cell strainer and purified by centrifugation 30min at 2400rpm in 40/80 Percoll (GE Healthcare) gradient. Lungs were finely minced and digested in complete RPMI supplemented with collagenase D (0.1mg/mL; Roche) and DNase I (20U/mL; Roche) for 1h at 37°C under gentle agitation. Sequentially, cells were passed through a 100μM cell strainer purified by centrifugation 30 minutes at 2400rpm in 40/80 Percoll (GE Healthcare) gradient. Spleen and mesenteric lymph node cell suspensions were obtained using 70μm strainers. Bone marrow cells were collected by either flushing or crushing bones and filtered using 70μm strainers. Erythrocytes from small intestine, colon, lung, spleen and bone marrow preparations were lysed with RBC lysis buffer (eBioscience). Leukocytes from blood were isolated by treatment with Ficoll (GE Healthcare).

### Flow cytometry analysis and cell sorting

For cytokine analysis *ex vivo*, cells were incubated with PMA (phorbol 12-myristate 13-acetate; 50ng/mL) and ionomycin (500ng/mL) (Sigma-Aldrich) in the presence of brefeldin A (eBioscience) for 4 hours prior to intracellular staining. Intracellular staining for cytokines and transcription factors analysis was performed using IC fixation and Staining Buffer Set (eBioscience). Cell sorting was performed using FACSFusion (BD Biosciences). Sorted populations were >95% pure. Flow cytometry analysis was performed on LSRFortessa X-20 (BD Biosciences). Data was analysed using FlowJo 8.8.7 software (Tree Star). Cell populations were gated in live cells, both for sorting and flow cytometry analysis.

### Cell populations

Cell populations were defined as: Bone marrow (BM) Common Lymphoid Progenitor (CLP): Lin^-^CD127^+^Flt3^+^Sca1^int^c-Kit^int^; BM Innate Lymphoid Cell Progenitor ILCP: Lin^-^CD127^+^Flt3^-^CD25^-^c-Kit^+^α4β7^high^; BM ILC1: CD45^+^Lin^-^ CD127^+^NK1.1^+^NKp46^+^CD49b^-^CD49a^+^; BM ILC2 progenitor (ILC2P): Lin^-^CD127^+^Flt3^-^ Sca1^+^CD25^+^; small intestine (SI) NK: CD45^+^Lin^-^NK1.1^+^NKp46^+^CD27^+^CD49b^+^CD127^-^ EOMES^+^ or CD45^+^Lin^-^NK1.1^+^NKp46^+^CD27^+^CD49b^+^CD127^-^; small intestine ILC1: CD45^+^Lin^-^NK1.1^+^NKp46^+^CD27^+^CD49b^-^CD127^+^Tbet^+^ or CD45^+^Lin^-^ NK1.1^+^NKp46^+^CD27^+^CD49b^-^CD127^+^; small intestine ILC2: CD45^+^Lin^-^ Thy1.2^+^KLRG1^+^GATA3^+^ or CD45^+^Lin^-^Thy1.2^+^KLRG1^+^Sca-1^+^CD25^+^; lamina propria, spleen, mesenteric lymph node and lung ILC3: CD45^+^Lin^-^Thy1.2^high^RORγt^+^ or CD45^+^Lin^-^Thy1.2^high^KLRG1^-^; ILC3 IL17^+^: CD45^+^Lin^-^Thy1.2^high^RORγt^+^IL17^+^; ILC3 IL22^+^: CD45^+^Lin^-^Thy1.2^high^RORγt^+^IL22^+^; for ILC3 subsets additional markers were employed: ILC3 NCR^-^CD4^-^: NKp46^-^CD4^-^; ILC3 LTi CD4^+^: NKp46^-^CD4^+^; ILC3 CCR6^-^NCR^-^: CCR6^-^ NKp46^-^; ILC3 LTi-like: CCR6^+^NKp46^-^; ILC3 NCR^+^: NKp46^+^; SI T helper (Th) 17 cells: CD45^+^Lin^+^Thy1.2^+^CD4^+^RORγt^+^; colon Tregs: CD45^+^CD3^+^Thy1.2^+^CD4^+^CD25^+^FOXP3^+^; colon Tregs RORγt^+^: CD45^+^CD3^+^Thy1.2^+^CD4^+^CD25^+^FOXP3^+^RORγt^+^. The lineage cocktail for BM, lung, small intestine lamina propria, spleen and mesenteric lymph nodes included CD3□, CD8α, CD19, B220, CD11c, CD11b, Ter119, Gr1, TCRβ, TCRγδ and NK1.1. For NK and ILC1 staining in the small intestine NK1.1 and CD11b were not added to the lineage cocktail.

### Antibody list

Cell suspensions were stained with: anti-CD45 (30-F11); anti-CD45.1 (A20); anti-CD45.2 (104); anti-CD11c (N418); anti-CD11b (Mi/70); anti-CD127 (IL7Rα; A7R34); anti-CD27(LG.7F9); anti-CD8α (53-6.7); anti-CD19 (eBio1D3); anti-CXCR4(L276F12); anti-NK1.1 (PK136); anti-CD3□ (eBio500A2); anti-TER119 (TER-119); anti-Gr1 (RB6-8C5); anti-CD4 (RM4-5); anti-CD25 (PC61); anti-CD117 (c-Kit; 2B8); anti-CD90.2 (Thy1.2; 53-2.1); anti-TCRβ (H57-595); anti-TCRγδ (GL3); anti-B220 (RA3-6B2); anti-KLRG1 (2F1/KLRG1); anti-Ly-6A/E (Sca1; D7); anti-CCR9 (CW-1.2); anti-IL-17 (TC11-18H10.1); anti-rat IgG1k isotype control (RTK2071); anti-streptavidin fluorochrome conjugates from Biolegend; anti-α4β7 (DATK32); anti-Flt3 (A2F10); anti-NKp46 (29A1.4); anti-CD49b (DX5); anti-Ki67 (SolA15); anti-rat IgG2ak isotype control (eBR2a); anti-IL-22 (1H8PWSR); anti-rat IgG1k isotype control (eBRG1); anti-EOMES (Dan11mag); anti-Tbet (eBio4B10); anti-FOPX3 (FJK-16s); anti-GATA3 (TWAJ); anti-CD16/CD32 (93); 7AAD viability dye from eBiosciences; anti-CD196 (CCR6; 140706) from BD Biosciences; anti-RORγt (Q31-378) and anti-mouse IgG2ak isotype control (G155-178) from BD Pharmingen. LIVE/DEAD Fixable Aqua Dead Cell Stain Kit was purchased from Invitrogen.

### Bone marrow transplantation

Bone marrow CD3^-^ cells were FACS sorted from *Arntl*^*fl*^, *Vav1*^Cre^*.Arntl*^*fl*^, *Rag1*^-/-^.*Arntl*^*fl*^, *Rag1*^-/-^.*Rorgt*^Cre^.*Arntl*^fl^, *Nr1d1*^+/+^, *Nr1d1*^-/-^ and C57BL/6 Ly5.1/Ly5.2 mice. 2×10^5^ sorted cells from *Arntl* deficient and competent wild type littermate controls, were intravenously injected in direct competition with a third-part wild type competitor (CD45.1/CD45.2), in a 1:1 ratio, into non-lethally irradiated NSG (150cGy) or *Rag2*^-/-^*Il2rg*^-/-^ (500cGy) mice (CD45.1). Recipients were analysed at 8 weeks post-transplantation.

### Quantitative RT-PCR

RNA from sorted cells was extracted using RNeasy micro kit (Qiagen) according to the manufacturer’s protocol. Liver, small intestine (ileum) and colon epithelium was collected for RNA extraction using Trizol (Invitrogen) and zirconia/silica beads (BioSpec) in a bead beater (MIDSCI). RNA concentration was determined using Nanodrop Spectrophotometer (Nanodrop Technologies). For TaqMan assays (Applied Biosystems) RNA was retro-transcribed using a High Capacity RNA-to-cDNA Kit (Applied Biosystems), followed by a pre-amplification PCR using TaqMan PreAmp Master Mix (Applied Biosystems). TaqMan Gene Expression Master Mix (Applied Biosystems) was used in real-time PCR. Real time PCR analysis was performed using StepOne and QuantStudio 5 Real-Time PCR systems (Applied Biosystems). *Hprt, Gapdh and Eef1a1* were used as housekeeping genes. When multiple endogenous controls were used, these were treated as a single population and the reference value calculated by arithmetic mean of their CT values. The mRNA analysis was performed as previously described^35^. Briefly, the comparative C_T_ method (2^-ΔCT^) in which ΔC_T (gene of interest)_ = C_T (gene of interest)_ -C_T (Housekeeping reference value)_ was employed. When fold change comparison between samples was required, the comparative ΔCT method (2^-ΔΔCT^) was applied.

### TaqMan Gene Expression Assays

TaqMan Gene Expression Assays (Applied Biosystems) were the following: *Hprt* Mm00446968_m1; *Gapdh* Mm99999915_g1; *Eef1a1* Mm01973893_g1; *Arntl* Mm00500223_m1; *Clock* Mm00455950_m1; *Nr1d1* Mm00520708_m; *Nr1d2* Mm01310356_g1; *Per1* Mm00501813_m1; *Per2* 00478113_m1; *Cry1* Mm00500223_m1; *Cry2* Mm01331539_m1; *Runx1* Mm01213404_m1; *Tox* Mm00455231_m1; *Rorgt* Mm01261022_m1; *Ahr* Mm00478932_m1; *Rora* Mm01173766_m1; *Ccr9* Mm02528165_s1; Reg3a Mm01181787_m1; *Reg3b* Mm00440616_g1; *Reg3g* Mm00441127_m1; *Muc1* Mm00449604_m1; *Muc2* Mm01276696_m1; *Muc3* Mm01207064_m1; *Muc13* Mm00495397_m1; *S100a8* Mm01276696_m1; *S100a9* Mm00656925_m1; *Epcam* Mm00493214_m1; *Apoe* Mm01307193_g1; *Cd36* Mm01307193_g1; *Fabp* Mm00444340_m1; *Fabp2* Mm00433188_m1; and *Scd1* Mm00772290_m1.

### Quantitative PCR analysis of bacteria in stools at the Phylum level

DNA from faecal pellets of female mice was isolated with ZR Fecal DNA MicroPrep(tm) (Zymo Research). Quantification of bacteria was determined from standard curves established by qPCR as previously described^2^. qPCRs were performed with NZY qPCR Green Master Mix (Nzytech) and different primer sets using a QuantStudio 5 Real-Time PCR System (Applied Biosystems) thermocycler. Samples were normalized to 16S rDNA and reported according to the 2^-ΔCT^ method. Primer sequences were: 16S rDNA, F-ACTCCTACGGGAGGCAGCAGT and R-ATTACCGCGGCTGCTGGC; Bacteroidetes, F-GAGAGGAAGGTCCCCCAC and R-CGCTACTTGGCTGGTTCAG; Proteobacteria, F-GGTTCTGAGAGGAGGTCCC and R-GCTGGCTCCCGTAGGAGT; Firmicutes, F-GGAGCATGTGGTTTAATTCGAAGCA and R-AGCTGACGACAACCATGCAC.

### *Citrobacter rodentium* infection

Infection with *Citrobacter rodentium* ICC180 (derived from DBS100 strain)^36^ was performed at ZT6 by gavage inoculation of 10^9^ colony forming units^36,37^. Acquisition and quantification of luciferase signal was performed in an IVIS Lumina III System (Perkin Elmer). Throughout infection, weight loss, diarrhoea and bloody stools were monitored daily.

### Colony forming unit measurement

Bacterial translocation was determined in the spleen, liver, and mesenteric lymph nodes, taking in account total bacteria and Luciferase^positive^ *C. rodentium*. Organs were harvested, weighted, and brought into suspension. Bacterial colony forming units (CFU) of organ samples were determined via serial dilutions on Luria Broth (LB) agar (Invitrogen) and MacConkey agar (Sigma-Aldrich). Colonies were counted after 2 days of culture at 37°C. Luciferase^positive^ *C. rodentium* was quantified in MacConkey agar plates using an IVIS Lumina III System (Perkin Elmer). CFU were determined per volume (mL) for each organ.

### Antibiotic and dexamethasone treatment

Pregnant females and new born mice were treated with streptomycin 5g/L, ampicillin 1g/L and colistin 1g/L (Sigma-Aldrich) into drinking water with 3% sucrose. Control mice were given 3% sucrose in drinking water as previously described^38^. Dexamethasone 21-phosphate disodium salt (200 μg) (Sigma) or PBS was injected intraperitoneally at ZT0. After 4, 8, 12 and 23 hours (ZT 4, 8, 12 and 23) mice were sacrificed and analysed.

### Chromatin immunoprecipitation (ChIP) assay

Enteric ILC3 from adult C57BL/6J mice were isolated by flow cytometry. Cells were fixed, cross-linked, lysed and chromosomal DNA-protein complex sonicated to generate DNA fragments ranging from 200-400 base pairs as previously described ^2^. DNA-protein complexes were immunoprecipitated, using LowCell# ChIP kit (Diagenode), with 1μg of antibody against ARNTL (Abcam) and IgG isotype control (Abcam). Immunoprecipitates were uncross-linked and analysed by quantitative PCR using primer pairs flanking ARNTL putative sites (E-boxes) in the *Ccr9* locus (determined by computational analysis using TFBS tools and Jaspar 2018). Results were normalized to input intensity and control IgG. Primer sequences were: A: F-CATTTCATAGCTTAGGCTGGCATGG; R-CTAGCTAACTGGTCTCAAAGTCCTC; B: F-GCCTCCCTTGTACTACCTGAAGC; R-TCCCAACACCAGGCCGAGTA; C: F-AGGGTCAATTTCTTAGGGCGACA; R-GCCAAGTGTTCGGTCCCAC; D: F-TCTGGCTTCTCACCATGACCACT; R-TCTAAGGCGTCACCACTGTTCTC, E: F-TTTGGGGAATCATCTTACAGCAGAG; R-ATTCATCCTGGCCCTTTCCTTCTTA; F: F-GCTCCACCTCATAGTTGTCTGG; R-CCATGAGCACGTGGAGAGAAAG; G: F-GGTCGAATACCGCGTGGGTT; R-CCCGGTAGAGGCTGCAAGAAA; H: F-AGGCAAATCTGGGCCTATCC; R-GGCCCAGTACAGAGGGGTCT; I: F-GGCTCAGGCTAGCAGGTCTC; R-TGTTTGGCCAGCATCCTCCA; J: F-ACTCAGAGGTGCTGTGACTCC; R-AGCTTTAGGACCACAATGGGCA.

### Food restriction (inverted feeding)

*Per1*^Venus^ mice fed during the night received food from 9pm to 9am (control group), whereas mice fed during the day had access to food from 9am to 9pm (inverted group). Food restriction was performed during 9 consecutive days as previously described^12^. For food restriction in constant darkness, *Per1*^Venus^ mice were housed in constant darkness with *ad libitum* access to food and water for 2 weeks. Then, access to food was restricted to the subjective day or night, for 12 days, in constant darkness.

### Inverted light-dark cycles

To induce changes in light regime, *Per1*^Venus^ mice were placed in ventilated, light-tight cabinets on a 12h light-dark cycle (Ternox). After acclimation, light cycles were changed for mice in the inverted group for 3 weeks to completely establish an inverse light cycle, while they remained the same for mice in the control group, as previously described^39^. For inverted light dark cycle experiments followed by constant darkness, after establishing an inverse light dark cycle, mice were transferred into constant darkness for 3 weeks.

### SCN lesions

Bilateral ablation of the SCN was performed in 9 to 12 week-old *Per1*^Venus^ males by electrolytic lesion using stereotaxic brain surgery, as described previously^15^. Mice were kept in deep anaesthesia using a mixture of isoflurane and oxygen (1-3% isoflurane at 1L/min). Surgeries were performed using a stereotaxic device (Kopf). After identification of bregma, a hole was drilled through which the lesion electrode was inserted into the brain. Electrodes were made by isolating a 0.25mm stainless steel insect pin with a heat shrink polyester tubing, except for 0.2mm at the tip. The electrode tip was aimed at the SCN, 0.3mm anterior to bregma, 0.20mm lateral to the midline, and 5.8mm ventral to the surface of the cortex, according to the Paxinos Mouse Brain Atlas, 2001. Bilateral SCN lesions were made by passing a 1mA current through the electrode for the duration of 6sec, in the left and right SCN separately. Sham lesioned mice underwent the same procedure, but no current was passed through the electrode. After surgery animals were housed individually under constant dark conditions with *ad libitum* food and water and were allowed to recover for 1 week before behavioural analysis. Successful SCN lesioned mice were selected based on MRI imaging, arrhythmic behaviour and histopathology analysis.

### Magnetic resonance imaging

Screening of SCN ablated mice was performed using a Bruker ICON scanner (Bruker, Karlsruhe, Germany). RARE (Rapid Acquisition with Refocused Echoes) sequence was used to acquire coronal, sagittal and axial slices (5 slices in each orientation) with the following parameters: RARE factor=8, TE=85ms, TR=2500ms, resolution = 156×156×500µm^3^ (30 averages). For high-quality images a 9.4T BioSpec scanner (Bruker, Karlsruhe, Germany) was employed. This operates with Paravision 6.0.1 software and interfaced with an Avance IIIHD console. Anatomical images (16 axial and 13 sagittal slices) were acquired using a RARE (Rapid Acquisition with Refocused Echoes) sequence with RARE factor=8, TE=36ms, TR=2200ms and resolution of 80×80×500µm^3^ (12 averages).

### Behavioural analysis

Sham-operated and SCN ablated mice were individually housed and after a 24h acclimation period their movement was recorded for 72h, starting at circadian time 7, in constant darkness, using the automated animal behaviour CleverSys system. Data were auto scored by the CleverSys software. Videos and scoring were visually validated. Circadian rhythmicity was evaluated by cosinor regression model^40,41^.

### Histopathology analysis

Mice infected with *Citrobacter rodentium* were sacrificed by CO_2_ narcosis, the gastrointestinal tract was isolated, and the full length of cecum and colon was collected and fixed in 10% neutral buffered formalin. Colon was trimmed in multiple transverse and cross-sections and cecum in one cross-section^42^, and all were processed for paraffin embedding. 3-4μm sections were stained with haematoxylin and eosin and lesions were scored by a pathologist blinded to experimental groups, according previously published criteria ^43-45^. Briefly, lesions were individually scored (0-4 increasing severity) for: 1-mucosal loss; 2-mucosal epithelial hyperplasia, 3-degree of inflammation, 4-extent of the section affected in any manner and 5-extent of the section affected in the most severe manner, as previously described^45^. Score was derived by summing the individual lesion and the extent scores. Mesenteric (mesocolic) inflammation was noted but not scored. Liver, gonadal and subcutaneous fat from *Arntl*^ΔRorgt^ were collected, fixed in 10% neutral buffered formalin, processed for paraffin embedding, sectioned into 3μm-thick sections and stained with haematoxylin and eosin. The presence of inflammatory infiltrates was analysed by a pathologist blinded to experimental groups. For the SCN lesions experiment, mice were sacrificed with CO_2_ narcosis, necropsy was performed, and brain was harvested and fixed in 4% PFA. Coronal section of 50µm thickness were performed in the vibratome (Leica VT1000 S), from 0.6 to −1.3 relatively to bregma, collected to Superfrost Plus slides (menzel-gläser) and let to dry overnight before Nissl staining. Stained slides were hydrated in distilled water for brief seconds and incubated in Cresyl Violet Stain solution (Sigma-Aldrich) for 30min. Slides were dehydrated in graded ethanol and mounted with CV Mount (Leica). Coronal sections were analysed for the presence/absence of SCN lesion (partial vs total ablation, unilateral vs bilateral), in a Leica DM200 microscope couple to a Leica MC170HD camera (Leica Microsystems, Wetzlar, Germany).

### Microscopy

Adult intestines from *Ret*^GFP^ mice were flushed with cold PBS (Gibco) and opened longitudinally. Mucus and epithelium were removed, intestines were fixed in 4% PFA (Sigma-Aldrich) at room temperature for 10 minutes and incubated in blocking/permeabilising buffer solution (PBS containing 2% BSA, 2% goat serum, 0.6% Triton X-100). Samples were cleared with benzyl alcohol-benzyl benzoate (Sigma-Aldrich) prior dehydration in methanol^18,46^. Whole-mount samples were incubated overnight or for 2 days at 4°C using the following antibodies: anti-Tyrosine hydroxylase (TH) (Pel-Freez Biologicals) and anti-GFP (Aves Labs). Alexa Fluor 488 goat anti-chicken and Alexa Fluor 568 goat anti-rabbit (Invitrogen) were used as secondary antibodies overnight at room temperature. For SCN imaging RFP^*ΔCamk2a*^ and RFP^*ΔRorgt*^ mice were anesthetized, perfused intracardially with PBS followed by 4% paraformaldehyde (pH 7.4, Sigma-Aldrich). The brains were removed and post-fixed for 24 hours in 4% paraformaldehyde and transferred to phosphate buffer. 50µm coronal sections were collected through the entire SCN using a Leica vibratome (VT1000s) into phosphate buffer and processed free-floating. Sections were incubated with neurotrace 500/525 (Invitrogen, N21480) diluted 1/200 and mounted using Mowiol. Samples were acquired on a Zeiss LSM710 confocal microscope using EC Plan-Neofluar 10x/0.30 M27, Plan Apochromat 20x/0.8 M27 and EC Plan-Neofluar 40x/1.30 objectives.

### RNA sequencing and data analysis

RNA was extracted and purified from sorted small intestinal lamina propria cells isolated at ZT5 and ZT23. RNA quality was assessed by an Agilent 2100 Bioanalyzer. SMART-SeqII (ultra-low input RNA) libraries were prepared using Nextera XT DNA sample preparation kit (Illumina). Sequencing was performed on an Illumina HiSeq4000 platform, PE100. Global quality of FASTQ files with raw RNA-seq reads was analysed using *fastqc* (ver 0.11.5) (https://www.bioinformatics.babraham.ac.uk/projects/fastqc/). *vast-tools*^47^ (version 2.0.0) aligning and read processing software was used for quantification of gene expression in read counts from FASTQ files using *VASTD-DB*^47^ transcript annotation for mouse genome assembly mm9. Only the 8443 genes with read count information in all 12 samples and an average greater than 1.25 reads/sample were considered informative enough for subsequent analyses. Preprocessing of read count data, namely transforming them to log2-counts per million (logCPM), was performed with *voom*^48^, included in the *Bioconductor*^49^ package *limma*^50^ (version 3.38.3) for the statistical software environment *R* (version 3.5.1). Linear models and empirical Bayes statistics were employed in differential gene expression analysis, using *limma*. For heatmaps, normalized RNAseq data was plotted using pheatmap (v1.0.10) R package (http://www.R-project.org/). Heatmap genes were clustered using Euclidean distance as metric. RNA–seq datasets analysed are publicly available in Gene Expression Omnibus repository with the accession number GSE135235.

### Statistics

Results are shown as mean ± s.e.m. Statistical analysis was performed using the GraphPad Prism software (version 6.01). Comparisons between two samples were performed using Mann-Whitney U test or unpaired Student t test. Two-way ANOVA analysis was used for multiple group comparisons, followed by Tukey post hoc test or Sidak’s multiple comparisons test. Circadian rhythmicity was evaluated by Cosinor regression model^40,41,51^, using cosinor (v1.1) R package. Single-component Cosinor fits one cosine curve by least squares to the data. The circadian expression Period was assumed to be 24h for all analysis and the significance of the circadian fit was assessed by a zero-amplitude test with 95% confidence. Single-component Cosinor yields estimates and defines standard errors with 95% confidence limits for Amplitude and Acrophase using Taylor’s series expansion^51^. The latter were compared using two-tailed Student t-test, where indicated. Results were considered significant at *P<0.05, **P<0.01, ***P<0.001.

---

**Extended Data Figure 1. Clock genes in progenitors and gut ILC. a**, Circadian clock gene expression in bone marrow innate lymphoid cell type 2 progenitor (ILC2P), small intestinal group 1 and 2 innate lymphoid cells (ILC1 and ILC2). ILC2P n=4; ILC1 n=3; ILC2 n=6. **b**, *Per1*^Venus^ MFI analysis in bone marrow innate lymphoid cell type 2 progenitor (ILC2P) and small intestine lamina propria ILC1 and ILC2. ILC2P n=6; ILC1 and ILC2 n=4. **c**, Circadian clock gene expression in bone marrow common lymphoid progenitor (CLP), innate lymphoid cell progenitors (ILCP) and innate lymphoid cell type 2 progenitor (ILC2P). n=3. **d**, Circadian clock gene expression in small intestine lamina propria group 1 and 2 innate lymphoid cells (ILC1 and ILC2). ILC1 n=3; ILC2 n=6. (b-d) White: light period; Grey: dark period. Data are representative of 3 independent experiments. (a,c,d) n represents biologically independent samples. (b) n represents biologically independent animals. Mean and error bars: s.e.m. (a) two-way ANOVA followed by Tukey’s multiple comparison test. P values are relative to differences for *Per1* expression in ILC1 and ILC2 when compared with ILC2P; (b,d) Cosinor regression was used to define circadian rhythmicity; *P<0.05; **P<0.01; ***P<0.001; ns not significant.

**Extended Data Figure 2. Circadian signals regulate gut ILC3. a**, Percentage of lamina propria group 3 innate lymphoid cells (ILC3); CD4^-^NCR^-^, LTi CD4^+^, and NCR^+^ ILC3 subsets. n=6. **b**, Cell numbers of lamina propria group 3 innate lymphoid cells CD4^-^NCR^-^, LTi CD4^+^, and NCR^+^ ILC3 subsets. n=6. **c**, Percentage of donor cells in group 2 innate lymphoid cells (ILC2), CD4^+^ T cells and Th17 subsets of mixed bone marrow chimeras. *Arntl*^*fl*^ n=6, *Arntl1*^Δ*Vav1*^ n=7. Data are representative of 3 independent experiments. (a-c) n represents biologically independent animals. Mean and error bars: s.e.m.. (a-c) two-tailed Mann-Whitney U test. **P<0.01; ns not significant.

**Extended Data Figure 3. Cell-intrinsic *Arntl* signals control intestinal ILC3. a**, Representative histogram of RFP expression in small intestine lamina propria ILC3. Representative of 3 independent analysis. **b**, Number of Peyer patches. *Arntl*^*fl*^ and *Arntl*^Δ*Vav1*^ n=9; *Arntl*^*fl*^ and *Arntl*^Δ*Rorgt*^ n=7. **c**, Percentage of small intestine lamina propria ILC3. n=5. **d**, Percentage of CCR6^-^NCR^-^, CCR6^+^ (LTi-like), and NCR^+^ ILC3 subsets. n=5. **e**, Percentage of intestinal lamina propria CD4^+^ T cells and helper (Th) 17 cells. n=5. **f**, Percentage and cell numbers of ILC3, regulatory T cells (Tregs) and RORγt expressing Tregs in the colonic lamina propria. n=6. **g**, Percentage of lamina propria ILC3, CCR6^-^NCR^-^, CCR6^+^ (LTi-like), and NCR^+^ ILC3 subsets. n= 3. **h**, Percentage of donor cells and cells numbers of intestinal ILC3 in mixed bone marrow chimeras. n=4. **i**, Percentage of donor cells in CCR6^-^NCR^-^, CCR6^+^ (LTi-like), and NCR^+^ ILC3 subsets of mixed bone marrow chimeras. n=4. **j**, Faecal Bacteroidetes and Firmicutes. *Arntl*^*fl*^ n=5; *Arntl*^Δ*Rorgt*^ n=6. **j**, White: light period; Grey: dark period. Data are representative of at least 3 independent experiments. (a-j) n represents biologically independent animals. Mean and error bars: s.e.m.. (b-f,h-i) two-tailed Mann-Whitney U test; (j) Cosinor regression was used to define circadian rhythmicity; (g) two-tailed unpaired Student t-test. *P<0.05; **P<0.01; ***P<0.001; ns not significant.

**Extended Data Figure 4. Impact of *Nr1d1* in intestinal ILC3. a**, Percentage and cell numbers of small intestine lamina propria ILC3. *N1d1*^+/+^ n=6; *N1d1*^-/-^ n=5. **b**, Percentage of CCR6^-^NCR^-^, CCR6^+^ (LTi-like), and NCR^+^ ILC3 subsets. *N1d1*^+/+^ n=6; *N1d1*^-/-^ n=5. **c**, Schematic representation of mixed bone marrow chimeras setting. **d**, Percentage and cell numbers of donor cells in mixed bone marrow chimeras. n=5. **e**, Percentage of donor cells in intestinal CCR6^-^NCR^-^, CCR6^+^ (LTi-like), and NCR^+^ ILC3 subsets of mixed bone marrow chimeras. n=5. Data are representative of at least 3 independent experiments. (a-e) n represents biologically independent animals. Mean and error bars: s.e.m.. (a,b,d,e) two-tailed Mann-Whitney U test; **P<0.01; ns not significant.

**Extended Data Figure 5. ILC3-autonomous ablation of *Arntl* impairs intestinal defence. a, b**, Histopathology of colon and cecum in *Rag1*^-/-^*.Arntl*^Δ*Rorgt*^ and *Rag1*^-/-^ *.Arntl*^*fl*^ littermate controls infected with *C. rodentium*. Pathological changes in colon and cecum of *Rag1*^-/-^*.Arntl*^Δ*Rorgt*^ mice included ulceration, loss of crypts and goblet cells, and inflammatory cell infiltration of the lamina propria by a granulocyte-rich population with a prominent and oedematous submucosa. Original magnification x4 (colon, scale bars: 500µm); ×1.25 (cecum, scale bars 250µm). **c**, Inflammation score in the cecum. n=5. **d**, Numbers of total ILC3; IL-17- and IL-22-producing ILC3. n=3. **e**, Whole-body imaging of *Rag1*^-/-^*.Arntl*^Δ*Rorgt*^ and their *Rag1*^-/-^*.Arntl*^*fl*^ littermate controls at day 7 after luciferase-expressing *C. rodentium* infection. **f**, MacConkey plates of liver cell suspensions from *Rag1*^-/-^*.Arntl*^Δ*Rorgt*^ and their *Rag1*^-/-^*.Arntl*^*fl*^ littermate controls at day 13 after *C. rodentium* infection. **g, h**, Total bacteria (left) and *C. rodentium* (right) translocation to the liver and mesenteric lymph nodes (mLN). n=4. **i**, Epithelial reactivity gene expression in the colon of *C. rodentium* infected *Rag1*^-/-^*.Arntl*^Δ*Rorgt*^ and their *Rag1*^-/-^ *.Arntl*^*fl*^ littermate controls. n=3. **j**, Weight loss in *C. rodentium*-infected mice. *Rag1*^-/-^ *.Arntl*^*fl*^ n=6; *Rag1*^-/-^*.Arntl*^Δ*Rorgt*^ n=7. **k, l, m**, Histopatholgy analysis of inflammatory infiltrates in the liver, gonadal and subcutaneous fat. Liver, scale bars: 250µm; gonadal and subcutaneous fat, scale bars: 100µm n=4. **n**, mouse total body weight. n=5. Data are representative of at least 3 independent experiments. (a-n) n represents biologically independent animals. Mean and error bars: s.e.m.. (c,g,h) two-tailed Mann-Whitney U test. (d,n) two-tailed unpaired Student t-test. (j) two-way ANOVA and Sidak’s test. *P<0.05; **P<0.01; ***P<0.001; ns not significant.

**Extended Data Figure 6. ILC3 proliferation, apoptosis and gut homing markers. a**, ILC3-related gene rhythmicity in the small intestinal lamina propria ILC3. n=4. **b**, RNA-seq analysis of lamina propria ILC3 at ZT23. n=3. **c**, Percentage of Ki67 expression in small intestine lamina propria ILC3. n=4. **d**, Percentage of donor cells in mixed bone marrow chimeras. n=4. **e**, Number of Lin^-^CD127^+^RORγt^+^ cells in the blood. n=4. **f**, Percentage of ILC3 in mesenteric lymph nodes (mLN). *Arntl*^*fl*^ n=6; *Arntl*^Δ*Rorgt*^ n=8. **g**, Diurnal expression of *Ccr9* transcripts in gut ILC3. n=4. **h, i**, Percentage of CCR9 expression in small intestinal lamina propria CCR6^-^NCR^-^, CCR6^+^ (LTi-like), and NCR^+^ ILC3 subsets, ILC2 and CD4^+^ T cells, n=3. **j**, Percentage of α4β7 expression in small intestine ILC1, ILC2 and CD4^+^ T cells, n=4. **k**, Percentage of CCR9 expression in gut ILC1, ILC2 and CD4 ^+^ T cells, n=4. **l, m**, Diurnal analysis of α4β7 and CXCR4 expression in small intestine ILC3. n=4. (a,e,g,l,m) White: light period; Grey: dark period. Mean and error bars: s.e.m.. (a,c-m) n represents biologically independent animals. (b) n represents biologically independent samples. (a,g) two-way ANOVA; (c,d,h-k) two-tailed Mann-Whitney U test; (e,l,m) Cosinor regression was used to define circadian rhythmicity. (f) two-tailed unpaired Student t-test. *P<0.05; **P<0.01; ***P<0.001; ns not significant.

**Extended Data Figure 7. Light entrains intestinal ILC3 circadian oscillations. a**, Inverted feeding regimens in constant darkness. *Per1*^Venus^ expression in gut ILC3. n=3. **b**, Circadian clock gene expression in hepatocytes of *Per1*^Venus^ mice in inverted feeding regimens. n=3. Acrophase mean and s.e.m: *Arntl*: control 0.4 ± 0.5, inverted 11.5 ± 0.2; *Per2*: control 15.2 ± 0.6, inverted 3.9 ± 0.5; *Nr1d1*: control 7.1 ± 0.6; inverted 18.5 ± 0.8. **c**, Opposing light-dark cycles. *Per1*^Venus^ in gut CCR6^-^NCR^-^, LTi CD4^+^, and NCR^+^ ILC3 subsets. n=3. Acrophase mean and s.e.m: CCR6^-^NCR^-^: control 14.5 ± 0.5, inverted 2.5 ± 0.5; LTi CD4^+^: control 14.5 ± 0.6, inverted 2.5 ± 0.4; NCR^+^: control 14.5 ± 0.6, inverted 2.5 ± 0.4. **d**, *Per1*^Venus^ MFI analysis of small intestine lamina propria ILC3 in mice maintained in constant darkness for 28 days. n=3. Data are representative of 3 independent experiments. (a-d) n represents biologically independent animals. Mean and error bars: s.e.m.. Cosinor regression. Standard errors with 95% confidence limits for amplitude (Amp) and acrophase (Acro) were extracted from the model and compared using two-tailed Student t-test. **P<0.01; ***P<0.001; ns not significant.

**Extended Data Figure 8. SCN-ablation shapes intestinal ILC3. a**, Circadian clock gene expression in small intestinal lamina propria ILC3. n=2-3. **b**, Magnetic resonance imaging of sham and SCN ablated (xSCN) *Per1*^Venus^ mice. Sagittal slices. White arrows: SCN ablation. **c**, Rhythms of animal locomotor activity. Total distance travelled in meters. **d**, Nissl staining of coronal brain sections. Top panel; scale bar: 1mm. Bottom panel; scale bar 250µm. (a-d) n represents biologically independent animals. Error bars show s.e.m.. Cosinor regression was used to define circadian rhythmicity; Cosine fitted curves are shown; Standard errors with 95% confidence limits for amplitude (Amp) and acrophase (Acro) were extracted from the model and compared using two-tailed Student t-test. *P<0.05; ns not significant.

**Extended Data Figure 9. Brain-tuned signals shape gut ILC3. a, b**, Confocal image of coronal brain sections showing neurotrace and RFP expression in the SCN. Scale bar: 100 µm. Representative of 3 independent analyses. **c**, Representative histogram of RFP expression in small intestine lamina propria ILC3. Representative of 3 independent analyses. **d**, *Per1* expression in small intestinal lamina propria ILC3. n=3. **e**, Percentage of small intestine lamina propria ILC3. n=3. **f**, Number of small intestine lamina propria ILC3. n=3. **g, h**, Epithelial reactivity gene expression in the intestinal epithelium. n=3. **i**, Rhythms of faecal Bacteroidetes and Firmicutes. *Arntl*^*fl*^ n=4, *Arntl*^Δ*Camk2a*^ n=3. (a-i) n represents biologically independent animals. Mean and error bars: s.e.m.. (d,g-i) Cosinor regression was used to define circadian rhythmicity; Cosine fitted curves are shown. (e) two-way ANOVA and Sidak’s test; (f) two-tailed unpaired Student t-test; *P<0.05; **P<0.0;1; ns not significant.

**Extended Data Figure 10. Impact of *Nr3c1* and *Adrb2* deficiency in gut ILC3. a**, *Per1*^Venus^ MFI analysis of lamina propria ILC3 after dexamethasone administration. n= 3. **b**, Percentage and cell numbers of small intestine ILC3. n=3. **c**, Percentage of lamina propria CCR6^-^NCR^-^, CCR6^+^ (LTi-like), and NCR^+^ ILC3 subsets. n=3. **d**, Tyrosine hydroxylase (TH) expressing neurons (red) and RET positive ILC3 (green) in cryptopatches. Scale bars: 40 μm. Representative of 3 independent analyses. **e**, Normalized expression of *Adrb*1, *Adrb2* and *Adrb3* in CCR6^-^NCR^-^, CCR6^+^, and NCR^+^ ILC3 subsets. **f**, Percentage and cell numbers of gut ILC3 in *Adrb2*^Δ*Il7ra*^ mice and their littermate controls. n=6. **g**, Percentage of lamina propria CCR6^-^NCR^-^, CCR6^+^ (LTi-like), and NCR^+^ ILC3 subsets in *Adrb2*^Δ*Il7ra*^ mice and their littermate controls. n=6. **h**, Percentage and cell numbers of gut ILC3 in *Adrb2*^Δ*Rorgt*^ mice and their littermate controls. *Adrb2*^fl^ n=3; *Adrb2*^Δ*Rorgt*^ n=4. **i**, Percentage of lamina propria CCR6^-^NCR^-^, CCR6^+^ (LTi-like), and NCR^+^ ILC3 subsets in *Adrb2*^Δ*Rorgt*^ mice and their littermate controls. *Adrb2*^fl^ n=3; *Adrb2*^Δ*Rorgt*^ n=4. **j**, light cues and brain-tuned circuits shape gut ILC3 homeostasis. Arrhythmic ILC3 impact intestinal homeostasis, epithelial reactivity, microbiota, enteric defence, and the host lipid metabolism. Thus, ILC3 integrate local and systemic entraining cues in a distinct hierarchic manner, establishing an organismal circuitry that is an essential link between diurnal light signals, brain cues, intestinal ILC3 and host homeostasis. (a-d,f-i) n represents biologically independent animals. (a) White: light period; Grey: dark period. Mean and error bars: s.e.m.. (a) two-way ANOVA and Sidak’s test. (b,c,f-i) two-tailed Mann-Whitney U test. *P<0.05;***P<0.001; ns not significant.

## Notes

https://www.ncbi.nlm.nih.gov/geo/query/acc.cgi?acc=GSE135235

